# A large-scale transcriptome-wide association study (TWAS) of ten blood cell phenotypes reveals complexities of TWAS fine-mapping

**DOI:** 10.1101/2021.02.23.432444

**Authors:** Amanda L Tapia, Bryce T Rowland, Jonathan D Rosen, Michael Preuss, Kris Young, Misa Graff, Hélène Choquet, David J Couper, Steve Buyske, Stephanie A Bien, Eric Jorgenson, Charles Kooperberg, Ruth J.F. Loos, Alanna C Morrison, Kari E North, Bing Yu, Alexander P Reiner, Yun Li, Laura M Raffield

## Abstract

Hematological measures are important intermediate clinical phenotypes for many acute and chronic diseases. Hematological measures are highly heritable, and although genome-wide association studies (GWAS) have identified thousands of loci containing trait-associated variants, the causal genes underlying these associations are often uncertain. To better understand the underlying genetic regulatory mechanisms, we performed a transcriptome-wide association study (TWAS) using PrediXcan to systematically investigate the association between genetically-predicted gene expression and hematological measures in 54,542 individuals of European ancestry from the Genetic Epidemiology Research on Adult Health and Aging (GERA) cohort. We found 239 significant gene-trait associations with hematological measures. Among this set of 239 associations, we replicated 71 at *p* < 0.05 with same direction of effect for the blood cell trait in a meta-analysis of TWAS results consisting of up to 35,900 European ancestry individuals from the Women’s Health Initiative (WHI), the Atherosclerosis Risk in Communities Study (ARIC), and BioMe Biobank. We further attempted to refine this list of candidate genes by performing conditional analyses, adjusting for individual variants previously associated with these hematological measures, and performed further fine-mapping of TWAS loci. To assist with the interpretation of TWAS findings, we designed an R Shiny application to interactively visualize TWAS results, one genomic locus at a time, by integrating our TWAS results with additional genetic data sources (GWAS, TWAS from other gene expression reference panels, conditional analyses, known GWAS variants, etc.). Our results and R Shiny application highlight frequently overlooked challenges with TWAS and illustrate the complexity of TWAS fine-mapping efforts.

**Author Summary:** Transcriptome-wide association studies (TWAS) have shown great promise in furthering our understanding of the genetic regulatory mechanisms underlying complex trait variation. However, interpreting TWAS results can be incredibly complex, especially in large-scale analyses where hundreds of signals appear throughout the genome, with multiple genes often identified in a single chromosomal region. Our research demonstrates this complexity through real data examples from our analysis of hematological traits, and we provide a useful web application to visualize TWAS results in a broadly approachable format. Together, our results and web application illustrate the importance of interpreting TWAS studies in context and highlight the need to carefully examine results in a region-wide context to draw reasonable conclusions and formulate mechanistic hypotheses.

## Introduction

Hematological measures (red cell, white cell, and platelet traits) have a critical role in oxygen transport, immunity, infection, thrombosis, and hemostasis and are associated with many acute and chronic diseases, including autoimmunity, asthma, cardiovascular disease, and COVID-19 [1–5]. Genome-wide association studies (GWAS) have identified thousands of loci containing such trait-associated variants, and previous Mendelian randomization and phenome-wide association study analyses have highlighted the likely causal role of blood cell trait-associated genetic variants in a variety of disorders, including autoimmune conditions and coronary heart disease [1–3]. Unfortunately, these individual SNP-based GWAS make it difficult to identify regulatory variants with small effect sizes which in aggregate impact the same gene, even in very large sample sizes, and they identify regions of associated variants whose biological function is often not clear [6]. Thus, utilizing a gene-based method to aggregate the effect of multiple regulatory variants may increase the study power to identify novel trait-associated loci and elucidate mechanisms of biological function.

A transcriptome-wide association study (TWAS) is one gene-based method which systematically investigates the association between genetically predicted gene expression and phenotypes of interest [6–9]. Here, we report results from a large TWAS of hematological measures using the PrediXcan method [6] to analyze data from 54,542 individuals of European ancestry from the Genetic Epidemiology Research on Adult Health and Aging (GERA) cohort (our discovery data set) [10] [11]. Hematological phenotypes are particularly good candidates for TWAS analysis due to the availability of large RNA-sequencing datasets in a relevant tissue type, high heritability across traits, and the large number of known genetic associations, most with poorly understood mechanisms and target genes. We perform this analysis using whole blood RNA-sequencing in 922 individuals from the Depression Genes and Networks (DGN) [12] study as our primary reference panel. After association analysis of imputed gene transcript levels with hematological indices in GERA, we performed conditional analyses, adjusting for variants previously identified to affect hematological measures, to evaluate if TWAS-identified genes represented novel statistical signals or were primarily driven by variants known from GWAS [3]. These direct conditional analyses represent a major advantage of the use of individual level data for our TWAS analyses, since these conditional tests could not be performed as easily or accurately using summary statistic-based methods. We replicated our significant set of gene-trait associations in a meta-analyzed sample of TWAS results containing 18,100 individuals from the Women’s Health Initiative (WHI), 9,345 individuals from the Atherosclerosis Risk in Communities Study (ARIC), and 8,455 individuals from Mount Sinai Bio*Me* Biobank (BioMe), all of European ancestry (Supplementary Table 1). We also compared the TWAS results from the primary DGN reference panel to three additional reference panels (whole blood and Epstein-Barr virus (EBV) transformed lymphocytes from the Genotype-Tissue Expression (GTEx) Project [13], and monocytes from the Multi-Ethnic Study of Atherosclerosis (MESA)[14]); these are considered secondary reference panels due to their smaller sample sizes. These additional analyses helped us to determine if relevant tissues with smaller sample sizes support our primary TWAS findings with DGN.

We employ several strategies to improve our understanding and interpretation of complex genomic regions containing multiple TWAS-identified genes. First, we used FOCUS (fine-mapping of causal gene sets [15]) to seek to identify a set of causal genes within genomic loci containing multiple significant TWAS gene-trait associations. FOCUS is a software used to fine-map TWAS statistics at genomic risk regions, while accounting for linkage disequilibrium (LD) among variants and predicted expression correlation among genes at those risk regions. Second, we present a novel web-based tool for integrating and visualizing TWAS and GWAS results, as well as results from multiple expression reference datasets. Additionally, we discuss frequently overlooked challenges of TWAS interpretation, such as failure to consider the number of proximal genes which cannot be accurately imputed with a given reference panel, but which may still be influenced by variants identified in GWAS studies. Our results illustrate the complexity of TWAS fine-mapping efforts but do provide one resource for clarifying likely gene targets for blood cell trait-related genetic loci. However, consideration of additional annotation resources and TWAS limitations is necessary for confident identification of gene targets.

## Results

We applied the PrediXcan method to identify expression-trait associations using individual level genotype and phenotype data from the GERA non-Hispanic white ethnic group. Analyzed blood cell traits included platelet count (PLT), red blood cell counts (red blood cell count (RBC), hematocrit (HCT), hemoglobin (HGB), mean corpuscular volume (MCV), and red cell distribution width (RDW) indices), and white blood cell counts (white blood cell count (WBC), monocyte count (MONO), neutrophil count (NEUTRO), and lymphocyte count (LYMPH) indices). We used DGN whole blood expression panel weights from PredictDB (a database of weights provided by PrediXcan; see URLs) to predict gene expression levels in GERA. Among the 11,538 genes in the DGN expression panel, 11,438 genes were predicted in GERA and 51% of those genes achieved DGN model R2 > 0.05 (see Supplementary Table 2 for model R^2^ values for significant genes and Supplementary Table 3 for genes included in DGN but not predicted in GERA). The number of GERA variants used for prediction was equal to the number of variants included in the prediction model (i.e. complete variant matching) for 74% of the predicted genes; the remaining genes used fewer variants from GERA than were included in the prediction models. We tested each of the 11,438 predicted genes individually for association with each of the 10 blood cell traits (BCTs), resulting in a Bonferroni-corrected p-value threshold of *p* < 4.37 × 10^−7^. Through the subsequent study analyses, we will refer to these results as “marginal TWAS”.

Overall, we identified 295 statistically significant marginal TWAS associations (*p* < 4.37 × 10^−7^), with each BCT having at least one significant association (Supplementary Table 2). Among these, 47 marginal TWAS associations fell into the major histocompatibility complex (MHC) or HLA region (GRCh37; chr 6: 28,477,797 - 33,448,354) and were not considered in subsequent analyses (Supplementary Table 4); disentangling a set of causal genes within the MHC region is exceptionally difficult due to the highly polymorphic genetic loci and complex LD in the region. Another 9 significant associations included genes which contained only a single variant in the prediction model. These associations were also not included in subsequent analyses (Supplementary Table 4). The remaining 239 significant associations include genes predicted from 2 to 112 variants, with a median of 21 variants used in predictive models. Among this set of 239 associations, we replicated 71 at *p* < 0.05 with same direction of effect for the blood cell trait in TWAS meta-analysis.

### Conditional analysis of significant TWAS genes on known GWAS variants

To determine whether any of the 239 remaining significant gene-trait associations were novel signals, not driven by any previously reported genome-wide significant variant, we performed conditional analysis. Since we performed TWAS with individual level data, we conditioned the predicted gene expression value of each statistically significant marginal TWAS gene on the set of nearby (within ± 1 Mb of the gene) sentinel GWAS variants within the BCT category (RBC, WBC, PLT) from the largest current European ancestry focused GWAS for BCTs [3]. We found that 222 (93%) of all marginal TWAS significant associations were attenuated by known GWAS variants and became non-significant (*p* > 4.37 × 10^−7^) upon conditional analysis. Another 15 of the associations remained significant after conditional analysis, and the remaining 2 associations did not have GWAS reported variants within a 1 Mb window of the gene (Supplementary Table 5). We attempted to replicate these 17 conditionally significant findings in a TWAS meta-analysis which included up to 32,036 European ancestry individuals from 3 cohorts: ARIC, WHI, and BioMe. Two of the 17 conditionally significant gene-trait associations (*HIST1H2BO*-HGB and *HIST1H2BO*-RDW) included in the replication set met the stringent significance threshold (*p* < 2.94 × 10^−3^). However, *HIST1H2BO* is situated within 1 Mb of the MHC region already excluded above (GRCh37; chr 6: 28,477,797 - 33,448,354), with this signal potentially reflecting long range LD with the MHC region, and has poor model R^2^ = 0.016. Additionally, *OR2B6* associated with HGB, MCV, and RWD; *ZNF192* associated with HGB, MCV, and RDW; and *ZSCAN12* associated with MCV meet a more lenient significance threshold (*p* < 0.05). Yet, *OR2B6*, *ZNF192*, and *ZSCAN12* are also located on chromosome 6 within approximately 500kb of the MHC region and all have poor model R^2^ < 0.015. The remaining 7 gene-trait associations did not meet any replication criteria.

### Characterization of TWAS Identified loci

Previously reported GWAS sentinel variants were at least partly responsible for and attenuated 93% of the significant marginal TWAS signals. Thus, we next examined if TWAS aided fine-mapping and identification of regulatory mechanisms at these loci. To better contextualize if fine-mapping in GERA was consistent in additional cohorts, we also examined replication of TWAS significant genes in these known loci. The 239 marginal TWAS associations reside in 120 trait-specific, physically non-overlapping *cis* loci (i.e. the *cis* region of each locus is ± 1 Mb of the locus’s TWAS sentinel gene start and stop positions). Over half (57%) of these loci contain only a single significant gene, while another 19% contain two significant genes. The remaining 24% of non-overlapping loci contain three or more significant genes, with up to 11 significant genes at a single locus. These 120 loci contain 87 unique index genes (defined as the most significant TWAS gene within the locus). Most loci do not contain a TWAS gene that replicates in meta-analysis (67% total; that is 47% of all loci are single-gene loci that do not replicate plus 20% of all loci are multi-gene loci that do not contain any gene that replicates, even at a marginal level). Ten percent of all loci are single-gene loci that meet a marginal replication threshold (*p* < 0.05), and 20% of all loci are multi-gene loci that meet this marginal threshold for at least one gene at the locus. The remaining 4 loci (3%) contain multiple TWAS genes with at least one gene meeting a strict replication threshold (*p* < 2.09 × 10^−4^) (Supplementary Table 2). These include the following index gene-phenotype associations: *TRIM68*-MCV, *USP49*-MCV, *PSMD3*-NEUTRO, and *PSMD3*-WBC.

### TWAS fine-mapping

In order to facilitate TWAS fine-mapping and allow for better interpretation of whether a given TWAS-identified gene is truly likely to associate with BCT variation, or whether it is likely to be a spurious association due to correlation of expression with nearby genes or other factors, we created an R Shiny application to interactively visualize TWAS sentinel genes in context, one locus at a time. The application allows us to integrate multiple sources of information from our primary TWAS analysis, including gene expression prediction models, TWAS meta-analysis, TWAS using secondary reference panels (whole blood and EBV transformed lymphocytes from GTEx, and monocytes from MESA), GWAS analysis of all BCTs, and correlation among genetic variants (i.e. LD) and among predicted gene expression levels. We highlight several loci to demonstrate the utility of the application, showcase some of the challenges that arise when TWAS identifies multiple significant genes at a single locus, and illustrate some of the complexities that are inherent in TWAS fine-mapping. In the sections that follow, we feature TWAS genes that fall into loci with a low, intermediate, or high level of complexity. All the figures in the following sections originate from the R Shiny application (http://shiny.bios.unc.edu/gera-twas/), which could be readily adapted to future TWAS analyses for other complex traits.

#### *HK1* locus

The *HK1* gene is known to be associated with several red blood cell traits including hemoglobin, mean corpuscular volume, hematocrit, mean corpuscular hemoglobin, red blood count, and red blood cell distribution width in GWAS analyses [3] and is a Mendelian gene for hemolytic anemia [MIM 142600]. Our TWAS results confirm previously reported *HK1* GWAS associations with HCT and MCV (assigned based on nearest gene for lead GWAS variants). The marginal TWAS tests for association between *HK1* and HCT (*p* = 3.84 × 10^−8^) and MCV (*p* = 1.05 × 10^−7^) are statistically significant (Fig 1); associations are all but eliminated by conditional analysis on known GWAS sentinel variants (HCT *p* = 2.58 × 10^−1^; MCV p = 4.36 × 10^−2^); *HK1* with HCT replicates in meta-analysis (p = 4.63 × 10^−2^); and *HK1* is the most significant TWAS gene among only two other genes (*HKDC1* and *TSPAN15*) implicated by GWAS at these loci, with the other two genes showing no TWAS signal.

**Figure 1.**
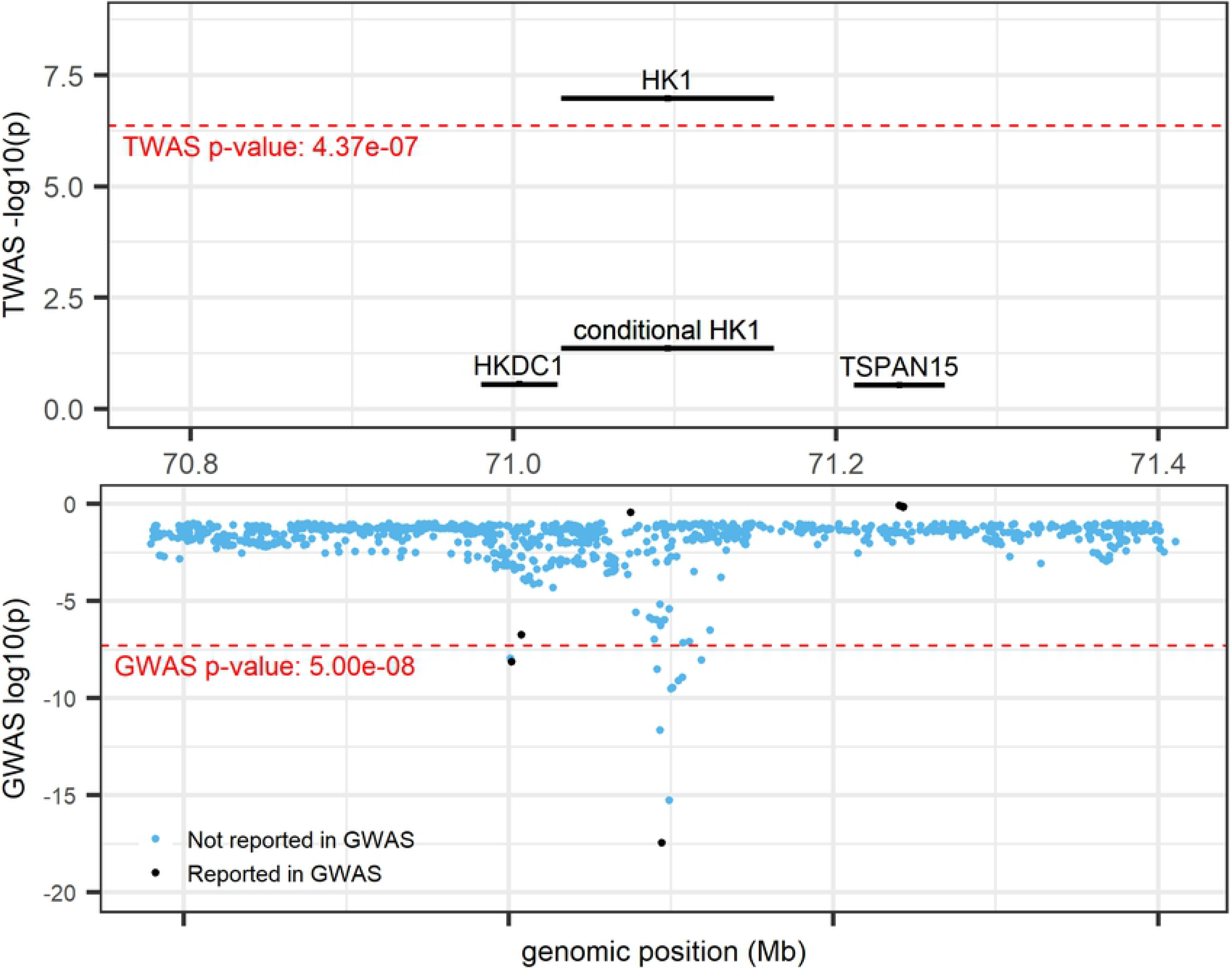
*HK1* locus (locus 60; chr 10: 70,029,740 - 72,161,638; trait = MCV) from R Shiny. TWAS results (top panel) and GWAS results (bottom panel). Marginal and conditional results for *HK1* are presented in the top panel. Black colored genes and variants denote those previously reported by UK Biobank and BCX GWAS [3], blue variants denote those not previously reported as UK Biobank and BCX GWAS sentinel variants.

#### *CREB5* locus

The marginal TWAS tests for association between *CREB5* and NEUTRO (*p* = 1.41 × 10^−12^) and WBC (*p* = 4.01 × 10^−10^) are the only TWAS significant associations at this locus (Fig 2A), and associations are essentially eliminated by conditional analysis on known GWAS sentinel variants (NEUTRO *p* = 9.04 × 10^−1^; WBC *p* =3.97 × 10^−1^). However, at this locus, both *CREB5* and *JAZF1* (TWAS NEUTRO *p* = 5.26 × 10^−3^, WBC *p* = 2.42 × 10^−3^) have previously been annotated as being the nearest and/or assigned gene for one or more GWAS sentinel variants. Predicted gene expression for *CREB5* and *JAZF1* are not highly correlated (r^2^ between 0.0 - 0.2) and appear to share only a single, non-significant predictive model variant (Fig 2B). *CREB5* and *JAZF1* both replicate at a lenient significance threshold for NEUTRO (*p* = 1.25 × 10^−2^, *p* = 8.98 × 10^−3^, respectively), and *CREB5* also replicates at a lenient threshold for WBC (*p* = 2.61 × 10^−2^) but *JAZF1*-WBC does not replicate (*p* = 0.11). Both genes appear in the GTEx whole blood (GWB) and MESA monocyte (MSA) secondary reference panels, but neither gene meets the significance threshold for either reference panel. Importantly, Human Protein Atlas [16] reports that *CREB5* is enhanced in blood and brain tissues and is specifically cell type enriched for NEUTRO [17]. *JAZF1* on the other hand has low tissue specificity [18]. Together the TWAS, GWAS, and Human Protein Atlas results point to *CREB5* as the most likely, and most biologically plausible, gene over *JAZF1* at this locus.

**Figure 2:**
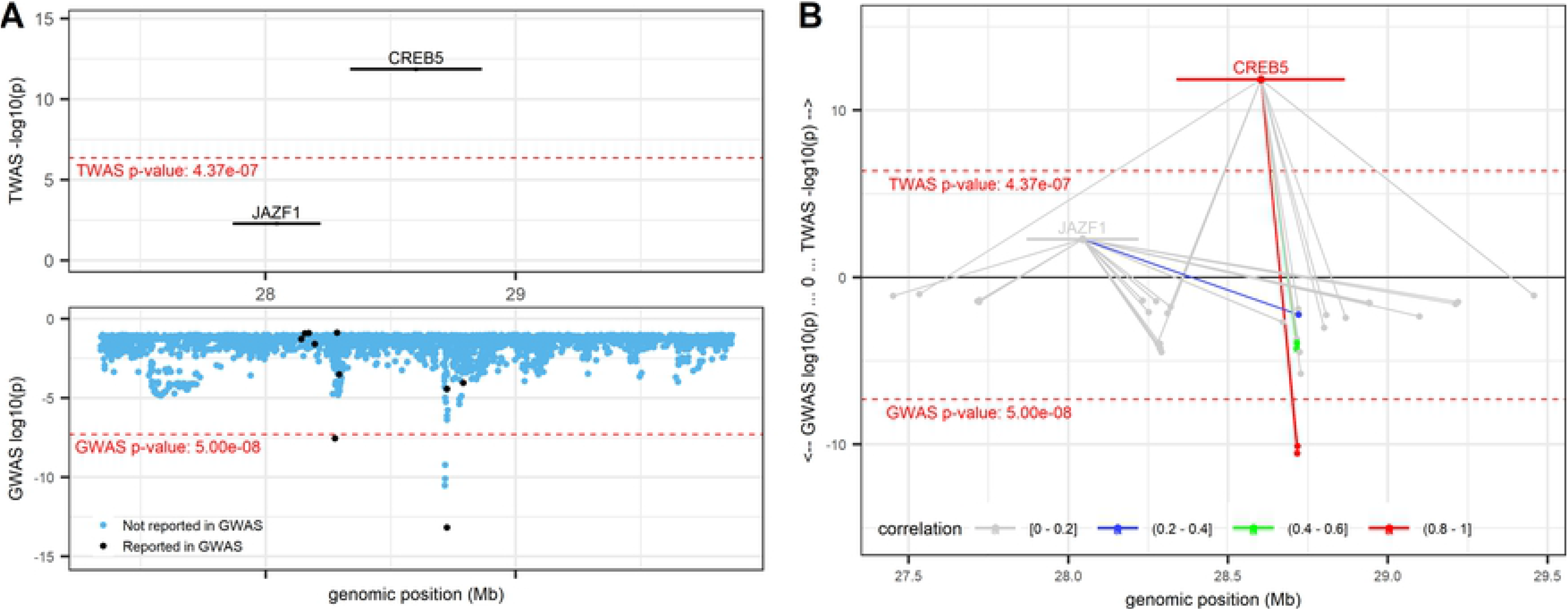
*CREB5* locus (locus 40; chr 7: 27,338,940 - 29,865,511; trait = NEUTRO) from R Shiny. TWAS results (top panels) and GWAS results (bottom panels). Marginal TWAS result displayed in (A), with Black colored genes and variants denoting those previously reported by GWAS, blue variants denote those not previously reported as GWAS sentinel variants. (B) Mirrored-Manhattan locus-zoom plot displaying genes connected to their predictive model variants. Color scale, increasing from light grey to red, indicates the predicted expression correlation (r^2^) between the index TWAS gene and all other genes in the locus and the LD between the index variant and all other variants in the locus.

#### *CD164* locus

The *CD164* gene is known to play a role in hematopoiesis [19, 20] and has been associated with several blood cell indices in GWAS analyses [3]. Our TWAS results prioritize *CD164* over other genes at the locus as being significantly associated with MCV (*p* = 2.54 × 10^−12^) and categorize it into a multi-gene locus along with *MICAL1* (*p* = 4.20 × 10^−7^) (Fig 3A). Conditional analysis on sentinel GWAS variants all but eliminates the TWAS signal for both *CD164* (*p* = 8.61 × 10^−2^) and *MICAL1* (*p* = 1.26 × 10^−1^). Interestingly, Fig 3B shows that *CD164* and *MICAL1* are not highly correlated in their predicted gene expression (r^2^: 0.2 - 0.4), and do not share any predictive model variants. We also note that both genes replicate in meta-analysis at a lenient threshold (*CD164 p* = 1.56 × 10^−3^; *MICAL1 p* = 7.06 × 10^−3^; Fig 3C). Additionally, while *MICAL1* is not available in secondary reference panels, *CD164* meets the TWAS significance threshold for its association with MCV in GTEx whole blood and MESA monocytes (GTEx *p* = 2.61 × 10^−9^ and MESA *p* = 4.09 × 10^−8^). Thus, the evidence at this locus suggests that expression of *CD164* and *MICAL1* are both reasonable candidates for being regulated by red cell-associated genetic variants although we note that Human Protein Atlas reports low tissue specificity for MICAL1 [21].

**Figure 3.**
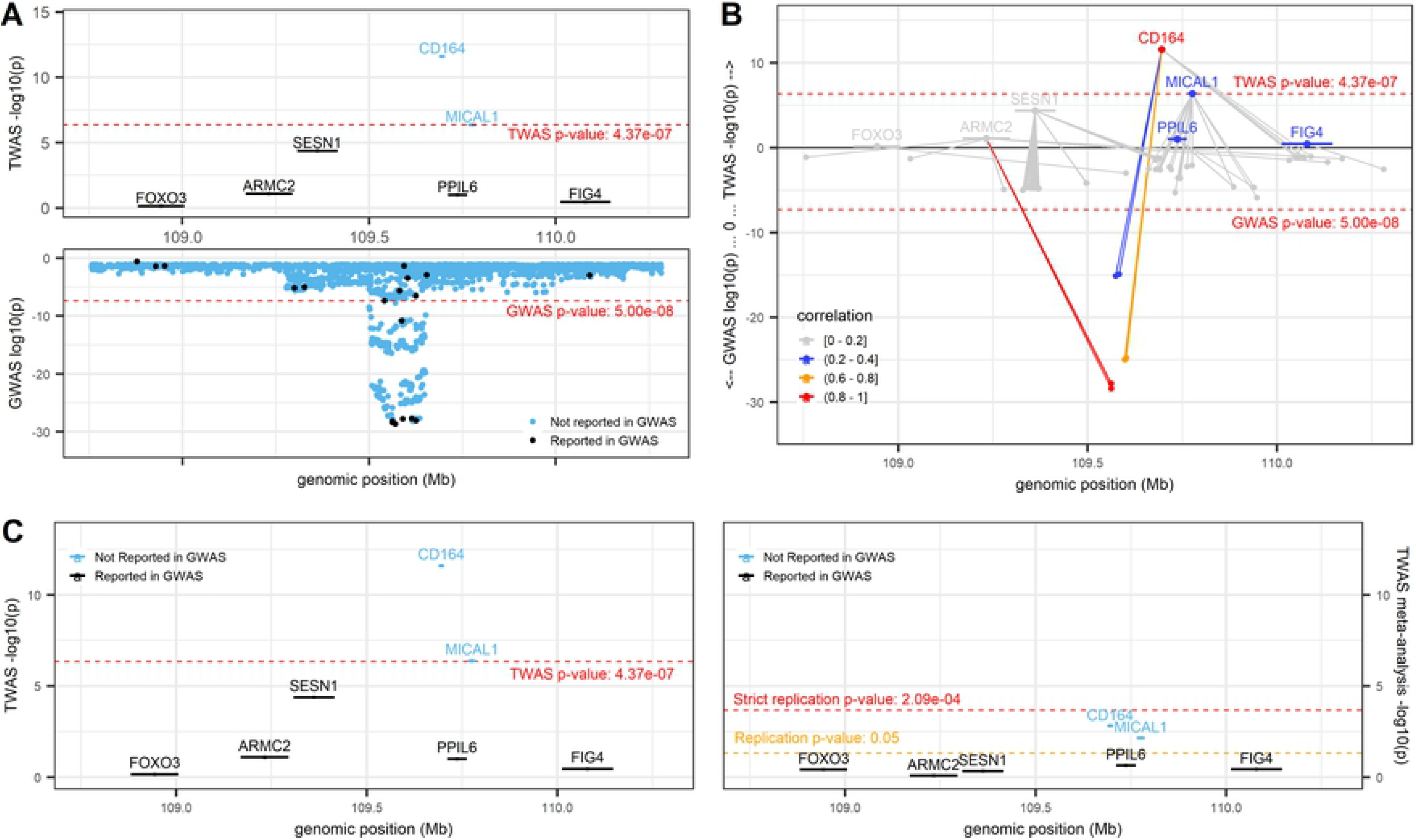
*CD164* locus (locus 36; chr 6: 108,687,717 - 110,703,762; trait = MCV) from R Shiny. (A) Marginal TWAS results in the top panel and GWAS results in the bottom panel. Black colored genes and variants denote those previously implicated by GWAS, and blue colored genes and variants denote those not previously implicated by GWAS. (B) Mirrored-Manhattan locus-zoom plot displaying genes connected to their predictive model variants. TWAS results in the top panel, GWAS results in the bottom panel. Color scale, increasing from light grey to red, indicates the predicted expression correlation (r^2^) between the index TWAS gene and all other genes in the locus and the LD between the index variant and all other variants in the locus. (C) Comparison of marginal TWAS (left panel) and TWAS meta-analysis (rightpanel). Black colored genes denote those previously implicated by GWAS sentinel variants, and blue genes denote those not previously implicated by GWAS sentinel variants.

#### *PSMD3* locus

The *PSMD3* locus contains a much higher level of complexity because it falls into a region containing many marginal TWAS genes, has a complex gene-gene correlation and LD pattern, and includes a combination of genes previously reported by GWAS as well as genes that have not been reported by GWAS. Thus, TWAS results do not clearly pinpoint the most likely causal gene. While *PSMD3* appears as the index TWAS gene associated with WBC (Fig 4A), 8 other genes are also TWAS significant at this locus. Five of those genes (*IKZF3, GSDMB, ORMDL3, MED24, CCR7*) replicate at a lenient significance threshold (*p* < 0.05), and *PSMD3* replicates at a more stringent threshold (*p* < 2.09 × 10^−4^) (Fig 4C). We see a complex network of shared model variants and correlation/LD patterns in Fig 4B, notably with *MED24* and *CCR7* (the next most significant genes at this locus) being only slightly correlated (r^2^ between 0.2 - 0.4) with *PSMD3*. The FOCUS fine-mapping results (Fig 4D) correspond to the TWAS results and indicate *PSMD3* and *MED24* as the most likely causal genes at the locus, each having posterior inclusion probabilities (PIPs) equal to 1.0. PIPs for all other genes at this locus, including *CCR7*, are less than 0.021 (Fig 4D).

**Figure 4.**
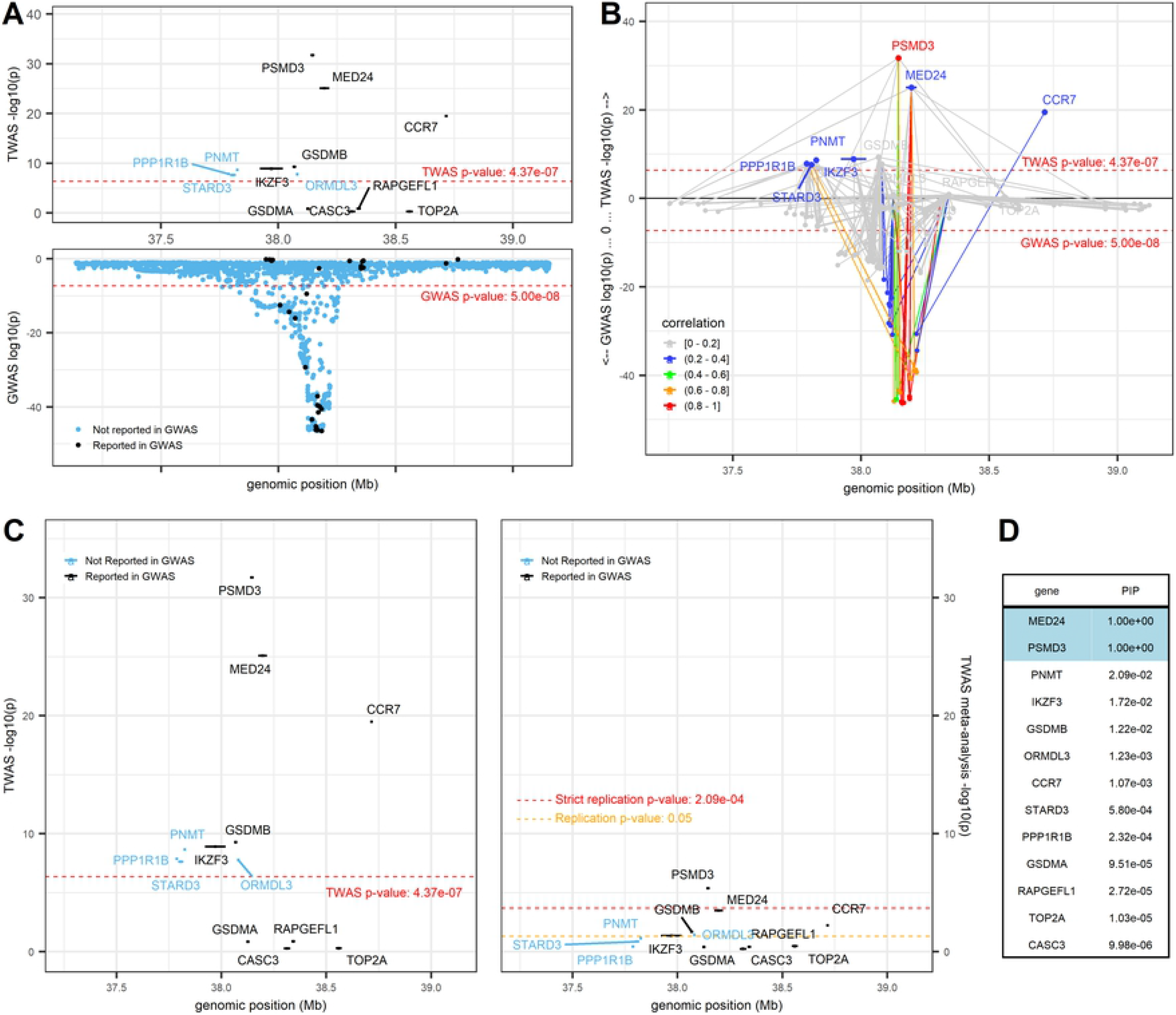
*PSMD3* locus (locus 101; chr 17: 37,137,050 - 39,154,213; trait = white blood cell count). (A) displays marginal TWAS results (top panel) and GWAS results (bottom panel), with genes and variants colored in blue and black to denote those not reported by GWAS and those reported by GWAS, respectively. (B) is a mirrored-Manhattan locus-zoom plot displaying genes connected to their predictive model variants with TWAS results (top panel) and GWAS results (bottom panel). Color scale, increasing from light grey to red, indicates the predicted expression correlation (r^2^) between the index TWAS gene and all other genes in the locus and the LD between the index variant and all other variants in the locus. (C) presents marginal TWAS results (left panel) and meta-analysis TWAS results (right panel), with genes colored in blue and black to denote those not reported by GWAS and those reported by GWAS, respectively. (D) displays the FOCUS posterior inclusion probabilities (PIPs) for each gene at this locus.

## Discussion

We performed a large-scale TWAS using PrediXcan on 54,542 GERA individuals of European ancestry and present compelling evidence that results from marginal TWAS analyses alone cannot illuminate causal genes at loci for complex traits. We also present results of conditional analysis of the TWAS significant genes. While 17 gene-trait associations did remain significant after conditional analysis or contained no known GWAS sentinel variants within a 1 Mb region of the gene, we found no substantive evidence from meta-analysis nor secondary reference panels to support these associations as novel discoveries for BCTs. Conditional analyses suggest that nearly all our TWAS findings are driven at least in part by GWAS sentinel variants from the largest recent European focused GWAS analysis for BCTs [3]. Most gene-trait associations no longer meet our TWAS-wide significance threshold after conditioning on GWAS sentinels. This is perhaps not surprising given the greater statistical power for this GWAS analysis, which was conducted in 563,085 participants. However, for 61 gene-trait associations (26%), some residual signal (*p* < 0.05) remains after conditioning on GWAS. For example, although *JAK2* is a well-known blood cell-associated signal from GWAS [3] and the Mendelian disease literature for platelet disorders [MIM 147796], its association with platelet count remained statistically significant after conditional analysis. Thus, our TWAS results suggest that there are likely additional regulatory variants at the *JAK2* locus which are not tagged by current GWAS single variants. Similarly, for other gene-trait associations retaining some residual significance after conditional analysis our results suggest that additional small-effect regulatory variants remain to be discovered for these genes which associate with blood cell indices. This illustrates the power advantages from aggregate tests like TWAS.

We grouped the 239 TWAS-wide significant gene-trait associations into 120 loci for fine-mapping. To effectively interpret these results, we introduce an R Shiny application which integrates TWAS and GWAS information into locus-specific, interactive visualizations which we use to assist with TWAS fine-mapping and interpretation. We show the utility of the R Shiny application in this endeavor by highlighting the varying levels of complexity at several TWAS loci and demonstrating where TWAS aligns with or provides advantages over GWAS. For example, the *HK1*-MCV locus shows a very simple genomic locus in which we find that TWAS confirms what we already know from GWAS. *HK1* is the predominant glucose phosphorylating enzyme in mammalian tissues that share a strict dependence on glucose utilization for their physiologic functions, such as brain, erythrocytes, platelets, lymphocytes, and fibroblasts [22], and coding variants in *HK1* are known to be associated with hemolytic anemia due to hexokinase deficiency [MIM 142600], providing a clear link to red blood cell related traits. The *CREB5* locus further demonstrates one of the advantages of TWAS over GWAS in that the TWAS results provide clarity regarding the likely causal gene at the locus. At this locus, *CREB5* and *JAZF1* have both been implicated by GWAS, again, likely assigned based on their physical proximity to the GWAS sentinel variant. However, *CREB5* shows a strong TWAS signal, replicates in the much smaller meta-analysis sample, and Human Protein Atlas provides clear evidence of enrichment in blood (specifically neutrophils), as compared to *JAZF1*, which shows low blood and immune cell specificity [16–18]. These results in aggregate support *CREB5* as the likely causal gene at this locus, even though *CREB5* may not be the closest gene in proximity to all sentinel GWAS variants within the region.

We specifically highlight the challenges, particularly at multi-gene loci, which should be taken into consideration when interpreting TWAS findings, including total and/or predicted expression correlation, shared predictive model variants, relevance of reference tissue panel, biological plausibility, etc. Furthermore, we demonstrate the importance of interpreting TWAS results in context. Although TWAS is useful for prioritizing candidate causal genes, researchers should guard against the hasty conclusion that the most significant gene is the only causal gene or even the most likely causal gene. For example, the conclusion at the *CD164* locus is not entirely clear based on TWAS results. While TWAS points to *CD164* as the causal gene, as does existing knowledge of the gene’s biological function, taking the full context of this locus into consideration, it is not out of the realm of possibilities that both *CD164* and *MICAL1* are causal at this locus. Furthermore, at the *PSMD3* locus we see potentially misleading TWAS results when marginal TWAS statistics are interpreted alone. The *PSMD3*-WBC association appears as the sentinel gene at this locus. However, there are several pieces of evidence which support other genes, including *CCR7*, as the most likely biologically plausible causal gene at the locus. First, *PSMD3* and *MED24* are ubiquitously expressed across tissues in Human Protein Atlas and have low tissue specificity and no immune cell specificity [16, 23, 24]. Additionally, results for these two genes differ slightly by reference panel. *MED24* appears in all secondary reference panels, but only exceeds the significance threshold in GWB and MSA. On the other hand, *PSMD3* only appears in the MSA reference panel and is not statistically significant in that panel. Second, *CCR7* is TWAS significant, it replicates at a lenient threshold in meta-analysis, and is enriched for expression in blood and lymphoid tissues, especially T-cells [25]. However, *CCR7* is not highly correlated with nor does it appear to share model variants with *PSMD3*, and the FOCUS results show a posterior inclusion probability of only 0.001. Finally, *CCR7* is known to be involved in the migration of neutrophils to lymph nodes [26]. While it is certainly possible at multi-gene TWAS loci for multiple genes at the locus to be contributing to trait regulation, it is also possible for spurious or non-relevant genes to be identified based on shared eQTLs across tissues which are not relevant to a given trait or correlation of gene expression.

Moreover, proximal genes which cannot be accurately imputed with a given reference panel, but which may still be influenced by variants identified by GWAS studies, must also be considered. For example, the gene colony-stimulating factor 3 (*CSF3*), which has a known key role in the production, differentiation, and function of granulocytes [MIM 138970], is also situated within the *PSMD3* locus. However, this gene has very low constitutive expression in whole blood [27], and it is not depicted in Fig 4 because a predictive model could not be fit for this gene in the DGN reference panel (likely due to very low expression); therefore, *CFS3* cannot be detected as a possible target gene at this locus (Supplementary Table 7 contains *CSF3*, along with other genes that have been assigned by one or more GWAS variants but are not included in DGN). This genomic region is extremely complex and highly pleiotropic, and any interpretation of this locus using TWAS results alone is likely to be overly simplistic. This complex locus shows the importance of considering statistical evidence from TWAS, GWAS, and FOCUS fine-mapping as well as trait biology in the interpretation of TWAS findings.

While we have used PrediXcan and pre-calculated PredictDB weights for our analysis, we note a limitation in doing so. The variants included in PredictDB were not always available in our analytical cohort (generally due to poor imputation quality), so some predictive models did not use all PredictDB weights. We note that 70% of our TWAS significant genes were predicted with complete variant matching (i.e. used all model variants) and 85% of TWAS significant genes used at least 90% of model variants; we have included this information in Supplementary Table 2 for transparency, and these details should be taken into account when interpreting TWAS results.

The cohorts that we have included in our TWAS meta-analysis also pose some limitations on our ability to replicate GERA TWAS sentinel genes. The smaller sample sizes of the meta-analyzed cohorts are likely the primary reason why GERA TWAS sentinel genes fail to replicate. Additionally, it may be the case that major contributing variants exhibit differential allele frequencies across cohorts; although this is less likely than in multi-ethnic analyses because all cohorts are of European ancestry, it could still contribute to poorer power for replication. Furthermore, differences in imputation quality across cohorts could also explain the failure to replicate TWAS sentinel genes in meta-analysis. Thus, future investigations using homogeneously imputed data are needed to ensure consistency in imputation quality across cohorts.

Although FOCUS, in some cases, helps to identify a set of the most likely causal genes at a locus, we have shown that it does not always provide enough evidence above and beyond TWAS to fully identify a putative causal gene set at a complex locus. Additionally, FOCUS performs a summary statistics-based TWAS method and then proceeds to fine-mapping the TWAS results from this method. However, we performed TWAS using PrediXcan, and thus, the fine-mapping results from FOCUS may not exactly match our PrediXcan TWAS results. In future, the FOCUS software could be extended to take pre-calculated TWAS results as input (using the TWAS method of the researcher’s choosing), bypassing the need to use GWAS summary statistics or to re-compute predicted gene expression. Our analysis is primarily conducted using whole blood TWAS weights only, with supplemental TWAS results available in our app for a few other blood-related tissues (whole blood and EBV transformed lymphocytes from GTEx and monocytes from MESA); we felt this was the most prudent approach to limit false positives and reduce needed multiple testing correction, versus an approach using TWAS weights in, for example, all GTEx tissues, particularly given the relatively large whole blood gene expression dataset available from DGN (N = 922). However, this choice could be inappropriate if the main relevant tissue at some blood cell-related loci is not in fact whole blood, and it limits our ability to use FOCUS fine-mapping to overcome choice of tissue for TWAS training. Joint/multiple tissue TWAS approaches such as UTMOST [9] and MR-JTI [8] could be employed in the future to assess the relevance of other tissues at blood-cell related loci.

In summary, we found that TWAS results enrich our understanding of GWAS, can help to explain trait variation, and are superior to merely selecting the nearest gene. We have shown that the gene, or genes, implicated in TWAS, in some cases, clearly overlap with what is known in GWAS and from prior knowledge of important genes in hematopoietic processes. However, while we show that TWAS may help in some cases to pinpoint likely causal genes, we emphasize the need for investigators to carefully interpret TWAS results alone, out of context. We introduce an R Shiny application and demonstrate its utility in assisting researchers in this endeavor by leveraging the TWAS and GWAS information available from the analytical cohort and interactively visualizing results one locus at a time. The results of this analysis are accessible online at http://shiny.bios.unc.edu/gera-twas/, and we also make the layout of this application available for others to import and analyze their own TWAS results in an R package called LocusXcanR, which is available on GitHub (https://github.com/amanda-tapia/LocusXcanR). Together with a clearer understanding of the relationship between TWAS and GWAS results and subject matter expertise, TWAS results can help us formulate mechanistic hypotheses for functional experimental validation.

## Materials and Methods

All cohorts are described individually below. We analyze 10 hematological phenotypes (platelet count, red blood cell count, hematocrit, hemoglobin, mean corpuscular volume, red cell distribution width, white blood cell count, monocyte count, neutrophil count, and lymphocyte count) across all cohorts.

### Ethics statement

We here performed secondary data analysis on deidentified data only (exempt research). All individual studies included were approved by relevant local institutional review boards, and participants provided written informed consent.

### PrediXcan method

PrediXcan [6] is a gene-based association test that prioritizes genes which are likely to be causal for the phenotype. It implements an elastic net-based method for selecting variants associated with gene expression in a given reference panel, and then uses those variants to predict gene expression in a cohort with only genotype data. We downloaded the PrediXcan software (see URLs) along with its prepackaged weights for gene expression data from PredictDB (see URLs). Weights for gene expression using RNA sequencing data were obtained from the Genotype-Tissue Expression project (version 7) [13] (whole blood, genes= 6208; and EBV transformed lymphocytes, genes=3000), Depression Genes and Networks [12] (whole blood, genes=11538, n=922), and Multi-Ethnic Study of Atherosclerosis (Europeans only, monocytes, genes=4647) [14]. Imputed genotypes for all cohorts were filtered for imputation quality based on R^2^ > 0.3; variants not meeting this threshold were excluded from the analysis. We use DGN as our primary reference panel for all TWAS analyses as it is the largest single whole blood RNA-seq dataset.

### Included cohorts

These TWAS analyses were limited to self-reported white or European ancestry participants, for easy comparability with the DGN European ancestry eQTL panel, including input of LD information into the R Shiny application (see R Shiny Methods), and with the largest single-ancestry blood cell trait GWAS.

#### Genetic Epidemiology Research on Adult Health and Aging (GERA)

The GERA cohort includes over 100,000 adults who are members of the Kaiser Permanente Medical Care Plan, Northern California Region (KPNC) and consented to research on the genetic and environmental factors that affect health and disease, linking together clinical data from electronic health records, survey data on demographic and behavioral factors, and environmental data with genetic data [10] [11]. By self-report, the GERA cohort is 81% White and 19% minority. Each GERA participant provided a saliva sample for extraction of DNA, which was conducted at KPNC using Oragene kits (DNA Genotek Inc., Ottawa, ON, Canada). DNA samples were genotyped at the Genomics Core Facility of UCSF. Genotyping was completed as previously described [10] using 4 different custom Affymetrix Axiom arrays with ethnic-specific content to increase genomic coverage. In addition to the QC protocols performed during genotyping, a total of six subjects, all female, were dropped due to sex non-agreement according to the Plink v1.07 --geno option and variants with more than 10% missingness were removed. Genotype data were phased without external reference using Eagle v2.3 and then imputed to 1000 Genomes Phase 3 v5 using Minimac3. Principal components analysis was used to characterize genetic structure in this European sample [11]. Hematological measures were extracted from medical records. In individuals with multiple measurements, the first visit with complete white blood cell differential (if any) was used for each participant. Otherwise, the first visit was used. In total, 54,542 non-Hispanic White individuals with hematological measures were included in the analysis.

#### Women’s Health Initiative (WHI)

WHI originally enrolled 161,808 women aged 50-79 between 1993 and 1998 at 40 centers across the US, including both a clinical trial (including three trials for hormone therapy, dietary modification, and calcium/vitamin D) and an observational study arm [28]. WHI recruited a socio-demographically diverse population representative of US women in this age range. Two WHI extension studies conducted additional follow-up on consenting women from 2005-2010 and 2010-2015. Genotyping was available on some WHI participants through the WHI SNP Health Association Resource (SHARe) resource, which used the Affymetrix 6.0 array (~906,600 SNPs, 946,000 copy number variation probes) and on other participants through the MEGA array [29]. Imputation and association analysis was performed separately in individuals with Affymetrix only, MEGA only, and both Affymetrix and MEGA data. For variants with both Affymetrix and MEGA genotypes available, MEGA genotypes were used. In total, 18,100 self-reported white women with hematological phenotypes were included. All WHI subcohorts were imputed to 1000 Genomes Phase 3. Six sub-cohorts from the WHI study were included in the meta-analysis and phenotypes were not collected uniformly across the cohorts. Sample size information for each phenotype is contained in Supplementary Table 8.

#### Atherosclerosis Risk in Communities Study (ARIC)

The ARIC study was initiated in 1987 and recruited participants age 45-64 years from 4 field centers (Forsyth County, NC; Jackson, MS; northwestern suburbs of Minneapolis, MN; Washington County, MD) to study cardiovascular disease and its risk factors [30], including the participants of self-reported European ancestry included here. Standardized physical examinations and interviewer-administered questionnaires were conducted at baseline (1987-89), three triennial follow-up examinations, a fifth examination in 2011-13, a sixth exam in 2016-2017 and a seventh exam in 2018-2019. Genotyping was performed through the CARe consortium Affymetrix 6.0 array [31]. ARIC European American genotype data were imputed to Haplotype Reference Consortium (HRC) [32]. In total, 9,345 European ancestry participants with hematological phenotypes were included in the analysis. All phenotypes were adjusted for study site, age, age squared, sex, and top ten PCs and were inverse normalized.

#### Mount Sinai Bio*Me* Biobank

The Mount Sinai Bio*Me* Biobank, founded in September 2007, is an ongoing, broadly consented EHR-linked bio- and data repository that enrolls participants non-selectively from the Mount Sinai Medical Center patient population. The Bio*Me* Biobank draws from a population of over 70,000 inpatient and 800,000 outpatient visits annually from over 30 broadly selected clinical sites of the Mount Sinai Medical Center (MSMC). As of September 2020, Bio*Me* has enrolled more than 50,000 patients that represent a broad racial, ethnic and socioeconomic diversity with a distinct and population-specific disease burden, characteristic of the communities served by Mount Sinai Hospital. Bio*Me* participants are predominantly of African (AA, 24%), Hispanic/Latino (HL, 35%), European (EA, 32%), and other ancestry (OA, 10%). All blood cell phenotype data, as well as demographic variables, were extracted from the patients’ EHRs. Genotyping was performed using the Illumina GSA array (~640.000 variants) and genotype data were imputed using the “1000G Phase 3 v5” reference panel. In total, 8,455 European ancestry participants with hematological phenotypes were included in the analysis. All phenotypes were adjusted for study site, age, age squared, sex, and top ten PCs and were inverse normalized. The Bio*Me* Biobank Program operates under a Mount Sinai Institutional Review Board-approved research protocol. All study participants provided written informed consent.

### Conditional analysis

For each statistically significant TWAS gene-trait association, the effect of predicted gene expression was conditioned on a set of previously reported GWAS sentinel variants from [3] meeting the following criteria: 1) the sentinel variant fell within a 1Mb region of the TWAS gene, 2) the trait with which the GWAS variant was associated matched the TWAS analytical trait or was within the same trait category as the analytical trait (platelets, red blood cell indices [hematocrit, hemoglobin, mean corpuscular volume, red blood cell count, red blood cell distribution width], white blood cell indices [white blood cell count, neutrophils, monocytes, lymphocytes]), and 3) the GWAS variant met an imputation quality threshold of R^2^ > 0.3. We used a modified version of the cpgen R package (see cpgen Methods) to perform the conditional analysis, accounting for a PLINK KING-robust kinship matrix [33], which used only genotyped variants and excluded those with minor allele frequency less than 5% and those missing more than 1% of SNPs.

### Meta-analysis and replication with ARIC, WHI, BioMe

In order to replicate the conditionally significant gene-trait association, we tested each association via a meta-analysis of the ARIC, WHI, and BioMe cohorts. As described above, PrediXcan was used to facilitate gene expression imputation and association in each cohort separately, and the meta-analysis association test was conducted using METAL [34].

Replication of the GERA significant gene-trait associations was performed using meta-analyzed TWAS results from ARIC, WHI, and BioMe. Nine gene-trait associations remained statistically significant after conditional analysis; for this set of genes, we defined a Bonferroni-corrected statistically significant replication threshold at p-value < 5.56 × 10^−3^. For the fine-mapping analysis, statistical significance of replicated genes was qualified based on two different thresholds -- a stringent threshold Bonferroni-corrected for all 239 statistically significant TWAS gene-trait associations at p-value < 2.09 × 10^−4^, and a more lenient threshold at p-value < 0.05.

### FOCUS

We used the Fine-mapping Of CaUsal gene Sets (FOCUS) [15] software to fine-map TWAS statistics at genomic risk regions. As input, we used GWAS summary data from GERA along with eQTL weights from PredictDB Depression Genes and Networks whole blood data, and the European LD reference panel from 1000 Genomes Phase 3. The software outputs a credible set of genes at each locus which can be used to explain observed genomic risk.

### Fine-mapping loci and locus categories

Fine-mapping loci refers to fine-mapping analysis of trait-specific genomic locations that contain, and are centered at, sentinel TWAS genes. That is, we take the set of trait-specific statistically significant TWAS genes, select the most significant gene in the set (the sentinel gene), and assign it to a locus along with any other statistically significant TWAS genes within a 1Mb window of the sentinel gene. We then select the next most significant TWAS gene which has not yet been assigned to a locus and continue in this fashion until all statistically significant TWAS genes have been assigned to a locus.

We define locus categories based on whether the locus contains a single gene or multiple genes and whether the locus replicates in TWAS meta-analysis at either a lenient or strict threshold. Thus, locus categories are defined as follows: 1=single gene locus, strict replication (p < 2.09E-04); 2=single gene locus, replication (p < 0.05); 3=single gene locus, no replication; 4=multi gene locus, strict replication (p < 2.09E-04); 5=multi gene locus, replication (p < 0.05); 6=multi gene locus, no replication.

### R Shiny

We use R’s convenient Shiny package (version 1.5.0, implemented in R 4.0.3) to produce the web application which displays our GERA TWAS results. The IdeogramTrack (https://rdrr.io/bioc/Gviz/man/IdeogramTrack-class.html) uses Genome Reference Consortium Human Build 37 (GRCh37) and UCSC cytogentic bands from http://hgdownload.cse.ucsc.edu/goldenpath/hg19/database/. All GERA TWAS results were produced using PrediXcan as described above.

For GWAS results, GERA phenotypes (log10 transformed for WBC subtypes) were based on inverse normalized residuals and adjusted for sex, age, age-squared, and the first 10 genetic principal components; analysis was done with Bolt LMM as implemented in rvtests [35], as used in the meta-analyses reported in [3]. We excluded those without a valid date of blood cell count measurement, with age < 18 years, or with discordant genotypic and phenotypic sex, as well as those with no blood cell trait data. The cohort also has longitudinal data; we preferentially selected the first visit with complete data for all measures. If no visit had complete data, we used the first available visit. We also excluded extreme blood cell measures: WBC>200×109 cells/L, HGB>20 g/dL, HCT>60%, and PLT>1000×109 cells/L. For WBC subtypes, we analyzed log10-transformed absolute counts obtained by multiplying relative counts with total WBC count. Custom Axiom arrays used for GERA genotyping have been previously described [36] [37], as has genotype calling with apt-probeset-genotype and generation of PCs using EIGENSOFT4.2 [11].

GERA conditional analysis results were produced using cpgen as described below. Known GWAS sentinel variants were obtained from [3]. Model weights and model variants were taken from our primary DGN reference panel from PredictDB (or secondary reference panels GWB, GTL, or MSA from PredictDB). Correlation of predicted expression among genes at the locus was calculated using R’s cor() function, and LD among variants was computed using plink --r2 (https://zzz.bwh.harvard.edu/plink/ld.shtml). We used ggplot2() to produce all figures, except the network visualization used visNetwork(). Tables were produced using the DT package (https://www.rdocumentation.org/packages/DT/versions/0.16).

### cpgen

We used the R package cpgen to perform conditional analysis of TWAS-significant genes, while accounting for a KING kinship matrix. However, cpgen is designed in such a way that it performs eigenvalue decomposition on the cohort sample for every function call. Since we had 239 TWAS-significant associations, this would have required eigenvalue decomposition on a sample of N ~ 55,000 for each of those 239 associations, a computationally burdensome calculation. Thus, we slightly modified the cpgen script. Specifically, we computed the eigenvalue decomposition on the GERA sample outside of the cpgen script (for each phenotype), and then subsequently loaded the appropriate eigenvectors and eigenvalues into the program, modifying the script so that it could take these eigenvectors and eigenvalues as input.

## Acknowledgements

The data and materials included in this report result from collaboration between multiple studies and organizations. The authors thank the staff and participants of GERA. The authors also thank the WHI investigators and staff for their dedication, and the study participants for making the program possible. A listing of WHI investigators can be found at: https://www.whi.org/researchers/Documents%20%20Write%20a%20Paper/WHI%20Investigator%20Long%20List.pdf. The authors thank the staff and participants of the ARIC study for their important contributions. The authors also thank all participants in the Mount Sinai Biobank all our recruiters who have assisted and continue to assist in data collection and management and are grateful for the computational resources and staff expertise provided by Scientific Computing at the Icahn School of Medicine at Mount Sinai.

## URLs

cpgen: https://github.com/cheuerde/cpgen

FOCUS: https://github.com/bogdanlab/focus

Human Protein Atlas: https://www.proteinatlas.org/

LocusXcanR R package for R Shiny application: https://github.com/amanda-tapia/LocusXcanR

LocusXcanR R Shiny application for GERA results: http://shiny.bios.unc.edu/gera-twas/

METAL: https://genome.sph.umich.edu/wiki/METAL_Documentation

Online Mendelian Inheritance in Man (OMIM): https://www.omim.org/

PLINK KING kinship matrix: https://www.cog-genomics.org/plink/2.0/distance#make_king

PredictDB: http://predictdb.org/

PrediXcan: https://github.com/hakyimlab/PrediXcan

R Shiny: https://shiny.rstudio.com/

